# Movement data link phenotypic traits to individual fitness in a nocturnal predator

**DOI:** 10.1101/2022.08.26.505323

**Authors:** Paolo Becciu, Robin Séchaud, Kim Schalcher, Céline Plancherel, Alexandre Roulin

## Abstract

Recent biologging technology reveals hidden life and breeding strategies of nocturnal animals. Combining animal movement patterns with individual characteristics and landscape features can uncover meaningful behaviours that directly influence fitness. Consequently, defining the proximate mechanisms and adaptive value of the identified behaviours is of paramount importance. Breeding female barn owls (*Tyto alba*), a colour-polymorphic species, recurrently visit other nest boxes at night. We described and quantified this behaviour for the first time, linking it with possible drivers, and individual fitness. We GPS-equipped 178 breeding pairs of barn owls from 2016 to 2020 in western Switzerland during the chick rearing phase. We observed that 65% of breeding females tracked were (re)visiting nest boxes while still carrying out their first brood. We modelled their prospecting parameters as a function of partner-, individual- and brood-related variables, and found that female feather eumelanism predicted the emergence of prospecting behaviour (less melanic females are usually prospecting), while increasing male parental investment increased female exploratory efforts. Ultimately, females would revisit a nest more often if they had used it in the past and were more likely to lay a second clutch afterwards, consequently having higher annual fecundity than non-prospecting females. Despite these apparent immediate benefits, they did not fledge more chicks. We highlight how phenotypic traits can be related to movement patterns and individual fitness through biologging associated with long-term field monitoring.

## Introduction

Biologging is crucial for understanding patterns of movement-related animal behaviour and how they are directly linked to the dynamics and persistence of populations. This technology is allowing for the discovery, quantification, and analysis of many previously unknown or unquantifiable behaviours [1, 2]. Nowadays, with the accumulation of enormous quantities of biologging data we can finally answer long-standing questions in behavioural and movement ecology, such as how the internal state (including individual phenotypic characteristics), social interactions and abiotic environment can affect movement patterns and consequently individual fitness [3–5]. Recently, studies focused on how certain phenotypic characteristics or internal states could affect movement-related behaviours [6], or how the latter would affect individual fitness and/or survival [7–9], but very rarely have studies explored the link between individual phenotype, its interactions with the environment, and consequent fitness output [10, 11]. Movement-related behaviours with clear trajectorial patterns are currently effortlessly recognisable as related with foraging [12, 13], resource searching [14], and commuting [15, 16], but in other circumstances movement patterns require more attention and a better understanding of the social and environmental contexts [3]. A deeper study of movement patterns paired with environmental context-related information can help in hypothesising possible causes, mechanisms, and consequences of a newly identified behaviour [3, 17, 18]. Recently, post-tracking analytical tools are easing this process, helping to recognise and find repeated and consistent movement-related patterns across a large pool of individuals in order to identify a behaviour [19–21]. Identifying and analysing recursive movement patterns, such as repeated visits to specific locations like foraging patches, nests, or dens, is a powerful way to detect behavioural structure in movement trajectories [19, 20, 22]. In general, visiting potential nest sites one or multiple times can be highlighted by recursive movement and will probably be associated with prospecting for a suitable breeding site. Hence, for individuals prospecting behaviour is crucial for sampling spatial and non-spatial information to collect knowledge for future breeding-site-choice decisions [23–27]. If not too expensive in terms of time and energy, acquiring important information related to the breeding site is expected to enhance individual fitness [28, 29]. Usually, birds prospect for nesting sites in the pre-breeding period [24, 27], but it is not rare to find post-breeding prospectors [30–32], especially after unsuccessful breeding attempts [33, 34]. On the other hand, it is not uncommon for multiple-brooded species to have individuals prospecting while breeding, specifically towards the end of chick rearing [35]. By combining post-tracking tools and data from nest monitoring, we observed and identified an unexpected at-night movement-related behaviour by breeding female barn owls *(Tyto alba)*. Several individuals were visiting other nest sites during their chick-rearing phase. Performing such behaviour could redirect time and energy from chick rearing to an activity of which we do not know the benefits and related costs. Barn owls are a colour-polymorphic species breeding in nest boxes, with males more involved than females in chick provisioning and with both males and females often raising two broods per season [36]. In this study, we present the (re)visiting of surrounding nest-boxes as possible prospecting behaviour of breeding female barn owls in order to understand possible triggers and fitness-related consequences. We could quantify this prospecting behaviour from GPS-equipped wild barn owl breeding pairs in western Switzerland over five years in an intensive agricultural area [37]. Combining the high-frequency sampling rate with a behavioural classification method and movement recursive analysis we were able to characterize barn owl movement-related behaviour including visits to nest boxes and time spent there [19, 38]. We expected that female prospecting behaviour to be related to (a) individual factors (i.e., age, body condition, melanism, etc.), (b) partner characteristics (i.e., male’s age, home range size, chick’s provisioning rate, etc.) and (c) variables related to the ongoing clutch (i.e., brood size, chicks growth rate, laying date, etc.). Each of these variables could reasonably cause a long- or short-term behavioural change that would translate into different movement patterns (e.g., onset and amount of prospecting for nest boxes). In addition, we predict that (d) previous knowledge of the surrounding nesting boxes (e.g., prospecting female already nested there) and their success history (e.g., often occupied in previous years) could be predicting the probability of a nest box to be visited [26, 32]. We also expected that (e) prospecting behaviour predicts the possibility of re-nesting and starting a second clutch [37]. Finally, (f) performing this behaviour could predict an increase in female fecundity (if prediction ‘e’ is met), but not necessarily a higher number of fledglings or better-quality fledglings [37, 39]. While previous tracking studies on prospecting behaviour focussed on larger scale prospecting (e.g., colonies and possible breeding areas) [25, 40], often triggered by failed reproductive attempts [33, 34], mostly in seabird species, our study shows the mechanisms of different breeding strategies in a cosmopolitan and colour-polymorphic species, such as the barn owl, with deep implication to its individual fitness. Furthermore, this is the first evidence of prospecting (at specific nesting locations) with high-resolution tracking in a nocturnal land bird.

## Methods

### Study area and barn owl population monitoring

The study was carried out between 2016 and 2020 in an intensive agricultural landscape in western Switzerland where a wild population of barn owls breeds in nest boxes attached to barns [41]. For the first two weeks after hatching, the adult females stay almost entirely in the nest providing warmth to their offspring and distribute food brought by the male. After this period, both parents hunt small mammals, with the male being the main contributor to the chick provisioning [9, 36]. Data relative to breeding biology such as laying dates, number of eggs, nestlings and fledglings, body measurements, plumage pheomelanism (colour) and eumelanism (spottiness) assessment were taken as part of annual monitoring of the species for over 30 years [37].

### GPS tags, deployment, behaviour annotation and home range size

178 female barn owls and 122 male partners were tagged using GiPSy-5 GPS tags (2016-2017) and Axy-Trek GPS and accelerometer tags (2018-2020) (https://www.technosmart.eu/, Technos-mart, Italy). Movement data was processed for behaviour annotation and calculation of home range size following Séchaud et al. (2021, 2022)[9, 38]. Details about GPS deployment, behavioural categorization and calculation of home range size are shown in the Electronic Supplementary Materials.

### Identification of prospecting behaviour

We quantified the recursive movement of 171 female barn owls at nest boxes around their own with the `recurse R package` [19], using the function `getRecursionsAtLocations`. This function calculates number of visits at specific given locations with the option to include a buffer radius and a residence time threshold. We applied a circular buffer of 20 meters around nest boxes to control for GPS errors and two temporal thresholds of 1 and 600 minutes to have high resolution data on their time spent close to the visited nest using `calculateIntervalResidenceTime` (calculating the residence time during user-specified intervals in the radius around each location) and the number of different locations (nest boxes) visited per night. From the initial `recurse` object data, we calculated a series of parameters per night and per individual: overall visits to other nest boxes, number of revisits per nest box, number of nest boxes visited, median and sum time spent at the nest boxes. These parameters gave us an idea of the different expressions of the prospecting behaviour. For example, some females have many revisits but only to few nest boxes, while others visit many different nest boxes but only one time per box. We also created a binomial variable, called “prospecting [0,1]”, to simplify the behaviour. If the female visited at least a nest site other than her own it would account for 1, else 0. As response variable in our analyses, we used:

1. `prospecting [0,1]`: if the behaviour was expressed at least once [1] or never [0];
2. `mean visits per night`: number of total visits to nest boxes divided by the number of nights the female was tracked for, giving an idea of the motivation to (re)visit, regardless of the number of nest sites (re)visited;
3. `nest boxes visited`: number of nest boxes visited at least once, to account for the diversity of places visited;
4. `median time at nest sites per night`: this parameter relates to the information acquisition of the nest box visited.

For the continuous parameters (`mean visits per night, nest sites visited, median time at nest sites per night`) we also created a version without including the non-prospecting females (prospecting = 0), in order to analyse variation within the prospecting females (Figure 1).

**Figure 1:**
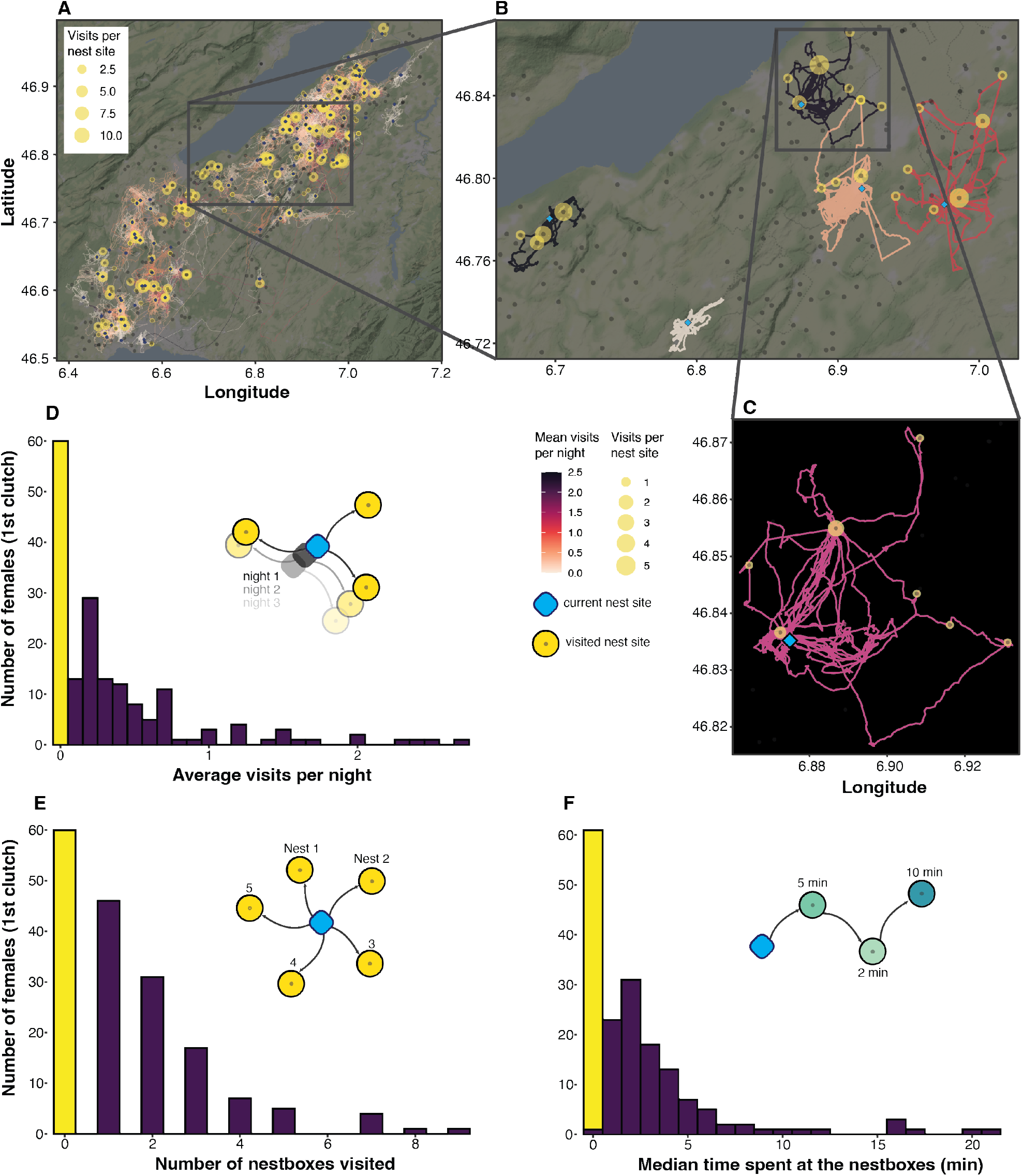
Schematic description and visualization of prospecting behaviour expressed by female barn owls. (A) Map of the study area with all tracked females considered for analysis (n = 171), highlighting average revisits per night (colour of the trajectories) and the number of visits per nest boxes (yellow circles). (B) Example of five female tracks with different degrees of prospecting behaviour or none (the lightest coloured track). (C) One track in detail, the colour of this track is for visualization purposes. (D-F) Histograms of the different parameters chosen to analyse the prospecting behaviour: in yellow females that did not visit any nest site except their own (n = 60), in violet the variation of the different prospecting parameters expressed by the females that visit at least one nest (n = 111).

### Explanatory variables

We divided the predictors into groups that match the expectations (a), (b) (c), (d), (e) and (f) in the Introduction:

a. Female characteristics that might influence their prospecting behaviour expression were considered. Daily mass variation, defined as the difference in body mass between tagging and tag retrieval divided by the number of days tag were deployed. Body condition as residuals of a linear regression between wing length and body mass [42] recorded at the moment of tagging. Age as factor (1^st^-year female or older than one year-since the exact year is not available for all individuals). Eumelanin expressed as number of spots per unit of space (count of spots in a 60×40 mm frame) on breast and belly, this factor can be related to different behavioural and breeding aspects mainly in female barn owls [43–45]. The latter variable is also correlated with spot diameter (Figure S1), hence we decided to use only the number of spots as less subjected to measurement error. Pheomelanin as breast and belly mean colour indexed from white (value of 8) to reddish-brown (value of 1), this trait was also found to be related with hunting behaviour and environmental factors [11, 46, 47].
b. In addition, we considered a set of variable relative to the female’s partner characteristics and behaviour. Body condition, age, eumelanin and pheomelanin as described above for the female parameters. We also added components of male hunting and movement behaviour such as home range size [**?**], provisioning rate per night defined as visits to their own nest by the males (see Séchaud et al., 2022 [**?**]), and proportion of time they spent perching, and therefore resting, at night as a proxy for their activity rate [38].
c. We also considered a group of variables relative to the ongoing breeding situation (during the tagging period). Brood size at the beginning of tag deployment as well as the loss of nestlings within the tagging period (brood size difference) as a proxy of pressure on parental food provisioning duties. We also considered the wing growth rate of the oldest chick in the brood during the tagging period to have an estimate of parental investment. Laying date (Julian date, as integer starting 1^st^ of January of every year) was also included to account for when in the breeding season they laid the first clutch eggs, usually the earlier the better for a second re-nesting chance in the same season [37].
d. The probability of visiting a specific nest box multiple times was placed in relationship to variables relative to the nest boxes themselves, such as: nest box usage in the previous 15 years (proportion), nest active in the current year (binomial), nest density (number of nest boxes active or not in a radius of 3.325 km around the current nest, radius corresponding to the 3^rd^ quartile of the visited nest boxes distances), distance (log-transformed) between breeding nest and visited nest boxes, number of breeding attempts by the female at visited nest boxes in the previous 15 years, and number of fledglings produced by any barn owl pair at visited nest boxes in the past 15 years, as a proxy for long-term nest box quality.
e. The explanatory variables used for predicting the probability of having a second clutch by the tracked female barn owls were laying date (for the aforementioned reason), brood size to account for the effort made in the first clutch, home range size of their partner in the first clutch as proxy of their quality [**?**], age of the female, final reproductive success of the first clutch expressed as fledglings produced divided the number of eggs laid. In addition, we included one of the four prospecting parameters after a variable selection among them (explained below).
f. Annual sum of eggs and fledglings were related to prospecting parameters as explanatory variables. Prior any statistical analyses we checked for correlations between the prospecting parameters and the number of available nest boxes in a range of 3.325, 5, 10, 20 km radius around the nest site of the tracked female. We did not find any strong correlation at any spatial scale hence we assumed that prospecting behaviour was not related to the availability of surrounding nest boxes (see Electronic Supplementary Materials). Also, female prospecting tracks are relative to their first clutch breeding event.

### Data analysis

In order to test our first three predictions (a), (b) and (c), concerning possible effects of partner-, brood- and female-related variables on the four prospecting parameters (`prospecting[0,1], mean visits per night, nest sites visited, median time at nest sites per night`), we created different subsets per prediction since the amount of missing data would differ between the sets of explanatory variables. Another series of subsets and models was also created excluding female barn owls that never visited a nest box other than their own (non-prospecting females) during the tracking period. Therefore, we built a series of (generalised) linear mixed models (GLMMs or LMMs) with different error distributions depending on the response variable: Binomial for `prospecting[0,1]`, Tweedie distribution for `mean visits per night and median time at nest sites per night`, Negative Binomial with quadratic parameterization for `nest sites visited`, Gaussian in models without non-prospecting females. The combination between the seven response variables and the three sets of predictors resulted in 21 (G)LMMs. To avoid multicollinearity issues, we chose the most biologically meaningful variable from pairwise Spearman rank correlation |*ρ*| > 0.6. This ensured that all the predictors in the GLMMs had a VIF (variance inflation factor) < 3 [50]. We then proceeded to select the optimal structure of the fixed component using a multi-model selection framework ranking the selected models according to the Akaike information criterion [51] using an automated stepwise model selection procedure in which models are fitted through repeated evaluation of modified calls extracted from the model containing all the meaningful variables, corrected for small sample sizes (AICc) [52]. Since no candidate model had a weight greater than 70%, parameters were estimated as the weighted averages of the set of models that represented 95% confidence weight [53] by averaging by the zero method [54]. To distinguish effects of predictors on response variables that differed from zero, 95% confidence intervals (C.I.) for each predictor were calculated. Furthermore, we used the Akaike weights (Σw_i_) [55] to assess the relative importance of the different variables (sum of Aikike weights, Σw_i_). To test if previous knowledge of surrounding nest sites could predict the probability of multiple visits to a specific nest site – prediction (d) – we ran a binomial GLMM with a response variable reflecting if a nest site was visited only once [0] or more than once [1]. To test prediction (e) that prospecting behaviour could possibly predict the probability of re-nesting (having a second clutch), we first set up a series of univariate binomial GLMMs using as binomial response variable if the female nested once or twice in the season (0 = one clutch, 1 = two clutches per year). We included as predictors the four prospecting parameters and then compared the four univariate models to choose the best predictor (model with lowest AICc). Using the best prospecting variable, we then built a more inclusive model including variables that could meaningfully predict re-nesting behaviour (as explained in the previous section). We followed the multi-model inference method described above to evaluate the most important predictors and their relationship with the probability of re-nesting a second time in prospecting and non-prospecting female barn owls. Finally, we tested possible long-term effects of prospecting behaviour – prediction (f) – evaluating the best prospecting predictor influencing the annual sum of eggs and fledglings by comparing univariate models as described above. We did not build a global model because the forces acting on annual fecundity and fitness are multiple, and our simplification is a starting point connecting the prospecting behaviour and two parameters of annual breeding output. We always included the year as a random effect, and we removed it from the model formula if the effect on estimate and C.I. calculation was negligible (variance was lower than 0.05 and estimates were the same with and without random factor). Prior to analysis, we centred and scaled the continuous predictors to mean zero and units of standard deviation (i.e., z-scores) to ensure comparability among variables. Also, we inspected GLMMs and LMMs residuals and considered the dispersion of the data [56] using a simulation-based approach to create readily interpretable scaled (quantile) residuals for fitted (G)LMMs with the package `DHARMa` [57]. Model fitting and multi-model inference were carried out in the statistical environment `R 4.1.0` [58] by the packages `glmmTMB` [59] and `MuMIn` [60], with `RStudio` [61] as graphic user interface. Descriptive statistics is reported as Mean *±* SD, unless specified otherwise.

### Ethics declaration

This study meets the legal requirements of capturing, handling, and attaching GPS devices to barn owls in Switzerland. All experimental protocols and methods were carried out in accordance with relevant guidelines and regulations of, and approved by, the Department of the Consumer and Veterinary Affairs, with legal authorizations: VD and FR 2844 and 3213; VD, FR and BE 3213 and 3571 (capture and ringing permissions from the Federal Office for the Environment).

## Results

The final dataset included 171 female barn owls – starting from 178 we filtered for tracks with a recording period of at least 3 nights (see Figure S2). 35% of females (n = 60) never visited another nest site other than their own (Figure 1D and 1E), while the remaining 65% (n = 111, min: 50% in 2017, max: 71% in 2016) visited another nest site at least once (see Figure 1 for a summary of their prospecting behaviour and its variability).

### Factors associated with prospecting behaviour

Generally, spottiness (number of melanic spots per unit of space) was found as key factor in our averaged models with prospecting (binomial), mean visits per night, number of nests visited, and median time spent at visited nest sites as response variables (Figure 2). Always scoring more than 0.9 variable predictive importance and having a negative effect, it was shown to be the most important variable. In the averaged model series of “prospecting [0,1]” the only variable that had an averaged coefficient with its 95% confidence interval not overlapping with zero was spottiness (*β* = −0.54, L.C.I. = −0.93, U.C.I. = −0.14, Σw_i_ = 0.95), while no other variable from models including brood-related and male-related variables was likely predicting the expression of the prospecting behaviour (first row in Figure 2, Table S1-3). On the other hand, when we considered the continuous response variables excluding the non-prospecting females – *n. nest sites visited, *median time at nest sites, *mean visits – and focussed only on the variation of the behaviour expression, we find several male-related variables as the most important predicting variables as well as the laying date as part of the brood-related factors (Figure 2, Table S7-9). Male body condition, home range size, and provisioning rate were the most important (Figure 2). We found that provisioning rate had positive effects on mean visits by their female mate (*β* = 0.20, L.C.I. = 0.02, U.C.I. = 0.39, Σw_i_ = 0.77) and median time spent at nest sites (*β* = 0.25, L.C.I. = 0.09, U.C.I. = 0.42, Σw_i_ = 0.96), meaning that the higher the attendance of the male at his own nest the more his partner female was visiting other(s) nest(s) and spending time there (Figure 2). Home range had moderate evidence of negative effect on number of nest sites visited (*β* = −0.17, L.C.I. = −0.31, U.C.I. = −0.02, Σw_i_ = 0.82) and mean visits (*β* = −0.19, L.C.I. = −0.37, U.C.I. = −0.001, Σw_i_ = 0.70), meaning that the smaller the male’s home range size the more the female prospected. However, this parameter was not found affecting their time spent at other nest sites. Male body condition showed contrasting results with moderate evidence of positive effect (*β* = 0.18, L.C.I. = 0.01, U.C.I. = 0.35, Σw_i_ = 0.72) on mean visits per night by the females, and a stronger negative effect (*β* = −0.24, L.C.I. = −0.4, U.C.I. = −0.08, Σw_i_ = 95) on median time spent at other nesting sites by their partners. Laying date was the only factor from the other two sets of variables that influenced one of the response variables. Specifically, the median time spent at other nest sites increased if the prospecting female started breeding (laying eggs) earlier (*β* = −0.21, L.C.I. = −0.39, U.C.I. = −0.02, Σw_i_ = 0.78). Similar results from male-related variables are shown for the same response variables but including the zeros from the non-prospecting females (n. nest sites visited, median time at nest sites, mean visits; Figure 2, Table S6). To conclude, the most important variable supported by strong evidence affecting all three response variables is the spottiness: more eumelanic or spotted females show less prospecting behaviour, consistently in all three variables (Figure 2, Table S5).

**Figure 2:**
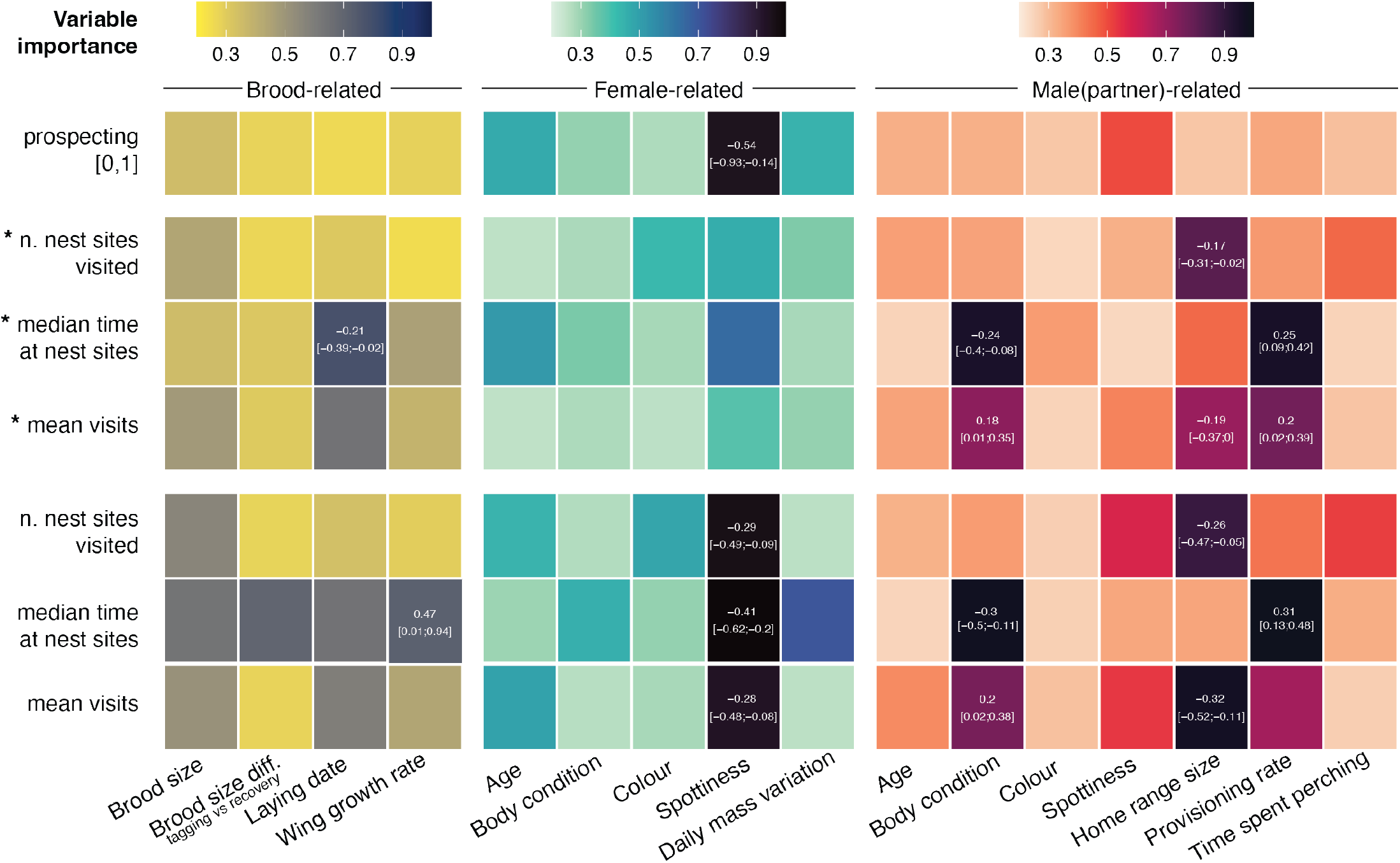
Summary of 21 averaged models (binomial, tweedie and negative binomial GLMMs) displaying the response variables on the y-axis and predictors on the x-axis. The response variables with a preceding “*” include only females that visit other nest sites during their tracking period. Tile colour refers to variable importance (see Methods for detailed explanation). The three different colour palettes highlight the three different set of predictors related with their breeding and brood in yellow to blue, the tracked female in blues, the male partner in red to dark purple. Estimate coefficients and 95% confidence intervals (C.I.) are reported when C.I. do not overlap 0, indicating moderate to strong evidence of the effects reported [48, 49].

### Drivers of multiple visits to a nest box

We tested if a series of parameters concerning nest sites in the area around the current breeding nest of the tracked females could influence their multiple revisitations. The binomial response variable summarises if the nest site is visited only once [0] or more than once [1]. The only factors affecting the probability of multiple visits to a specific nest box were previous nesting attempts by the tracked female in that specific nest box in the previous 15 years, and the distance to that nest box (Figure 3A, Table S10). Specifically, the probability of a nest to be visited more than once increased from roughly 25% to more than 50%, differing if the nest box visited was never previously used by the tracked female or there was at least one breeding attempt in the 15 years prior (Figure 3B). Other parameters concerning surrounding nest density, such as if the nest was currently occupied or not, how many times was occupied and mean fledged chicks in the previous 15 years were not found to affect the probability of multiple visits (Figure 3A).

**Figure 3:**
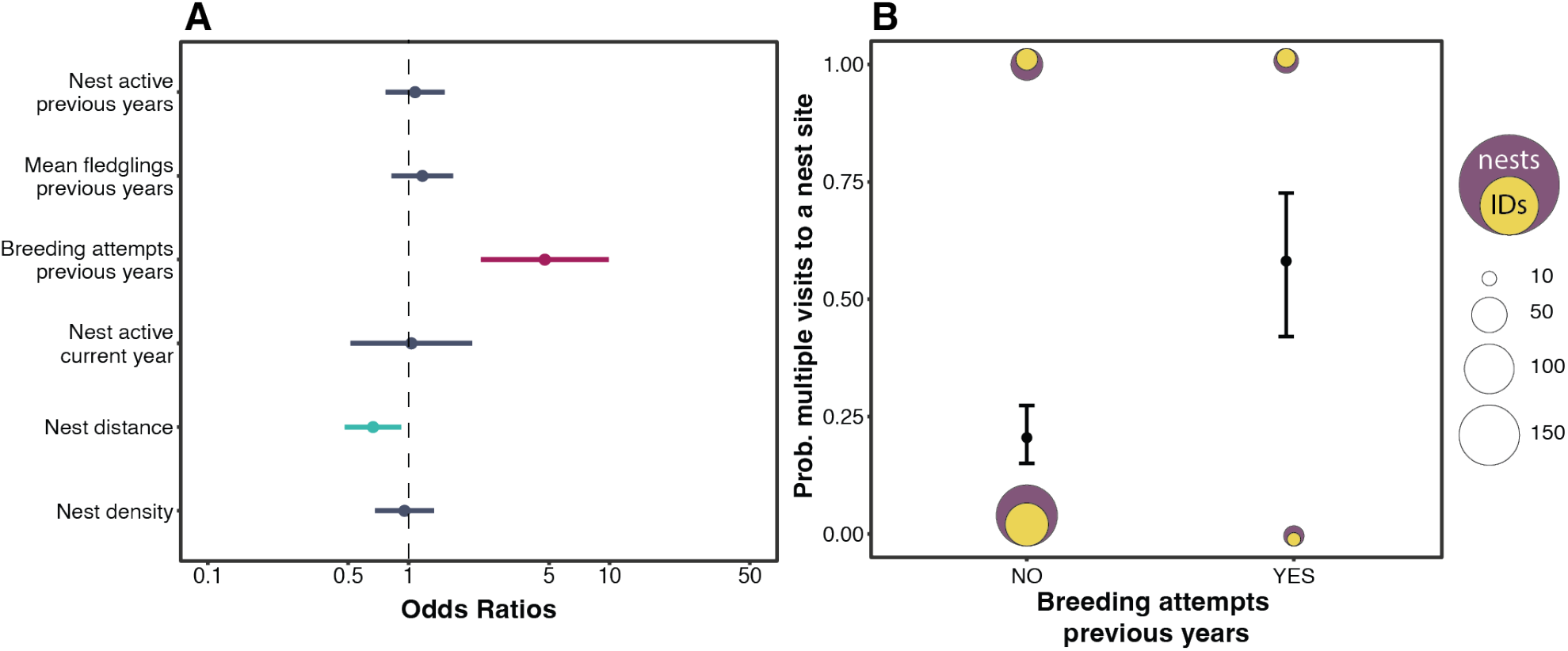
Binomial GLM summary of the results (estimated coefficient with 95% confidence interval ranges) on the probability of revisiting a nest site one or multiple times (A). Predicted effect (estimate with 95% confidence intervals) of the predictor with strong evidence of a positive effect given by breeding attempts in the revisited nest in previous years (B). Purple dots are nests visited and their size is related to the number of visits per category, yellow dots represent the number of individuals involved.

### Mean revisits predict probability of re-nesting

We found the mean visits per night was the best prospecting parameter predicting the probability of a second clutch (Figure 4A) with ΔAICc > 2 relative to the other univariate models (Table S11). We then included the mean visits per night in a more general binomial GLMM including brood size, age (of the female), home range size (of the male partner) and the proportion of fledglings relative to the number of eggs laid. From the global model we excluded the laying date since it is known to have the largest effect on re-nesting in this species (see [37]), and we confirm this finding in the Supplementary Material (Table S13, Figure S3). Our model showed moderate evidence of mean visits per night (Figure 4B, Table S12) with a positive predicting relative contribution on the probability of starting a second clutch (Figure 4C).

**Figure 4:**
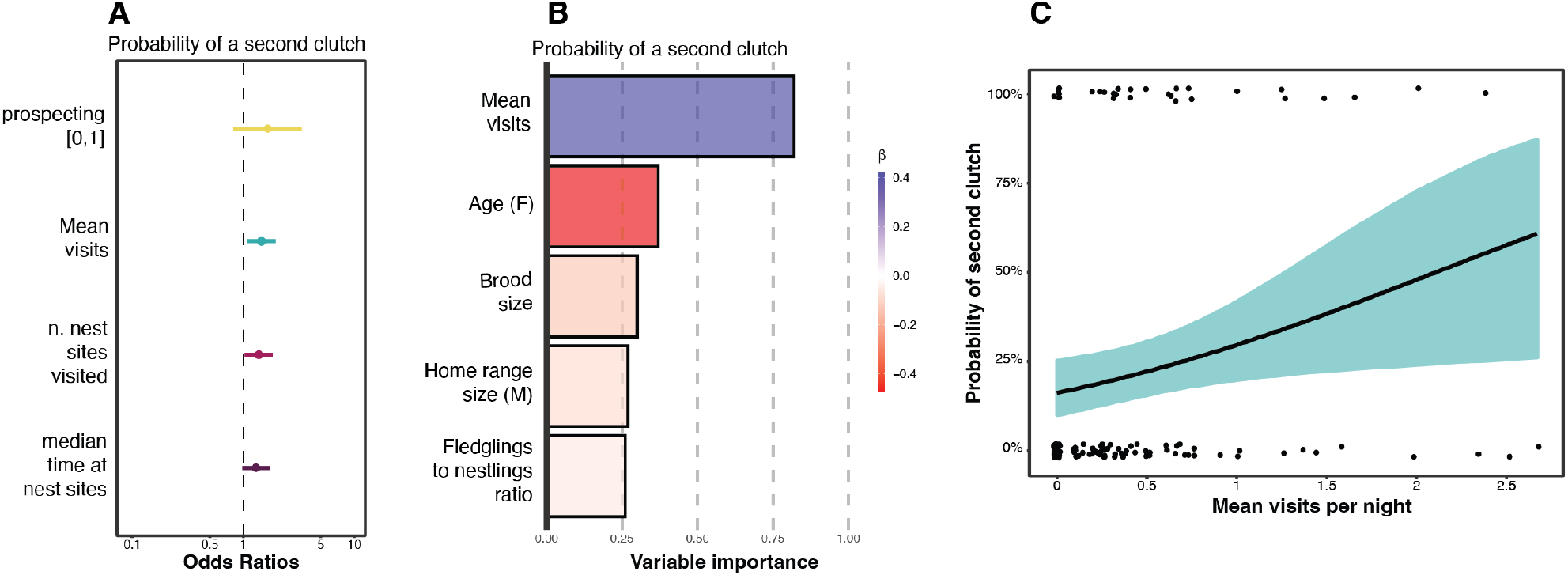
Set of univariate model results (estimated coefficient with 95% confidence interval ranges) with prospecting parameters as predictors of the possibility of having a second clutch by female barn owls (A). Summary of variable importance and model coefficients of averaged model (B), length of the bar is relative to variable importance (sum of Aikike weights – Σw_i_), colour is relative to the effect direction (red = negative, blue = positive) and its intensity is proportional to the effect strength. Predicted effect of mean visits per night (C) from averaged model in (B) on the probability of re-nesting for the second time in female barn owls, coloured area around the regression line is relative to 95% confidence intervals.

### Implications on annual reproductive fitness

As previously described, we ran a univariate model comparison between the prospecting parameters predicting female annual fecundity (sum of eggs per year) and female annual productivity (sum of fledglings per year). For the models with annual eggs as a response variable, the model comparisons highlighted the number of nest sites visited as the best predictor (AICc = 885.83, *β* = 0.084, U.C.I. = 0.139, L.C.I. = 0.028), followed by mean visits per night (AICc = 889.68, *β* = 0.061, U.C.I. = 0.118, L.C.I. = 0.005), the median time at nest sites (AICc = 890.06, *β* = 0.058, U.C.I. = 0.115, L.C.I. = 0.003) and the prospecting[0,1] (AICc = 891.23, *β* = 0.107, U.C.I. = 0.231, L.C.I. = −0.016) (Figure 5A, Table S14). We found that a female expressing the prospecting behaviour was more likely to have more eggs laid at the end of the breeding season (Figure 5B and 5C). On the contrary, no prospecting parameter seemed predict the sum of fledglings (Figure 5D, 5E, 5F, Table S15).

**Figure 5:**
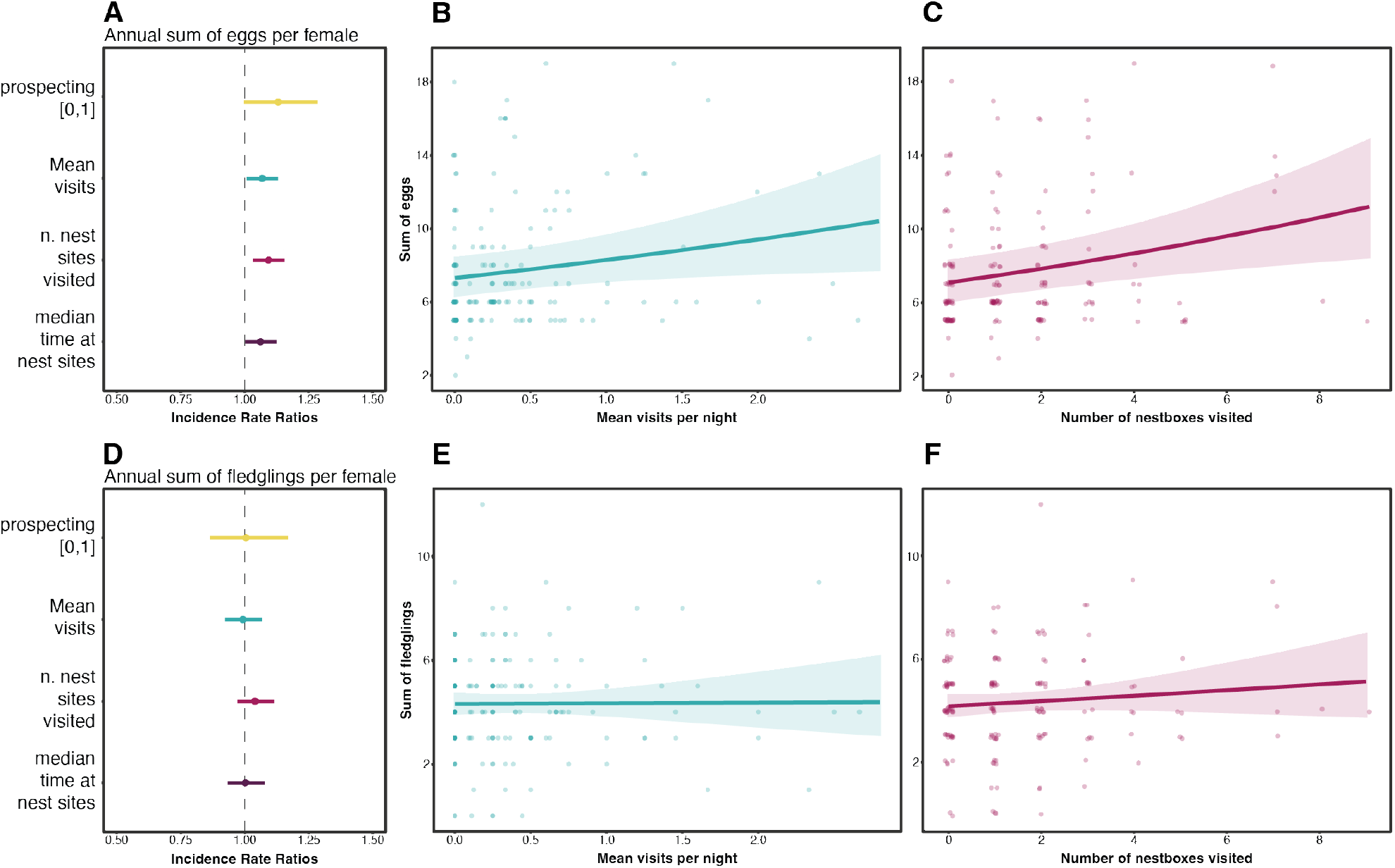
Set of univariate GLMM results (estimated coefficient with 95% confidence interval ranges) with prospecting parameters as predictors of annual sum of eggs (A) and annual sum of fledglings (D) produced by female barn owls. B, C, E and F panels represent the relationship (regression line and 95% confidence interval area) between two prospecting parameters and the annual sum of eggs and fledglings.

## Discussion

In this study, we described, quantified, and linked a previously unknown prospecting behaviour to possible causes and fitness consequences in breeding females of a nocturnal cosmopolitan predator, the barn owl. Prospecting for breeding sites is known to be a signal of breeding motivation that can be either triggered by external pressures, experience, or genetic predisposition [62, 63]. The variation linked to this strategy could also involve and affect parental investment and extra-pair activities, which we do not discuss in depth here. Also, we cannot exclude that female barn owls recorded as non-prospecting during the tracking period, did in fact prospect at other times. The emergence of female prospecting behaviour was found to be related to lack or reduced expression of eumelanin-based colouration (number of dark spots). In vertebrates, melanin-based coloration is often associated with variation in physiological and behavioural traits that probably stems from pleiotropic effects of genes regulating the synthesis of brown to black eumelanin [43]. Heavily spotted barn owl females were found to be more resistant to ectoparasites [64], to breed at earlier age [44], to be docile depending on their body condition [65] and preferable to males that had also a higher feeding rate [66]. Therefore, our finding regarding heavily spotted females being less explorative meaning less time invested in extra-clutch activities (i.e., prospecting for other nest boxes) appears in line with previous studies. Furthermore, these non-prospecting females had a similar annual reproductive fitness than prospecting ones, although the latter laid more eggs (increasing fecundity). A recent study found that barn owl fledglings with larger feather black spots reached farther distances when dispersing from their natal nest to their first breeding nest site [67]. This trait seems related with movement even at an early stage of barn owl life. Although this finding seems to contrast what we found, we should consider dispersal and prospecting while breeding as separate movement behaviours, since we cannot exclude that melanic females disperse more at an early stage of life (before their first breeding) and then do not prospect further during their following breeding seasons, and vice versa for less-melanic females. The variation of prospecting movement among females is likely predicted by their partner behaviour. Male provisioning rate positively affects the time spent at visited nest boxes and the mean visits per night, apparently allowing the female partner to wander more and be less tied to parental duties. On the contrary, smaller male home ranges triggered higher prospecting in terms of nests visited and mean visits per night. Home range sizes is probably related with higher quality areas and males having higher chances to raise more fledglings [9]. Finally, we found a contrasting effect of male body condition, having a stronger negative effect on time spent at other nest sites by prospecting females and a moderate positive effect on mean visits per night. On one hand, the first effect indicates that the more the female dedicates time to prospect, the more parental effort is expressed by the male that then pays the price with his body condition. The latter result, on the other hand, shows that males in better condition to be associated with higher mean visits per night, although the effect is very weak. Here, we show an association between male-associated behaviours and their partner’s behaviour, one of very few examples in the wider literature where individual movement-based behaviour has been shown to affect that of the respective partner [68]. In the future, we expect that with a growing collection of high-resolution biologging data, in avian and non-avian species, as well as an increasing theoretical and statistical framework on intra- and interspecific interactions [3, 5, 21, 69] these findings will be more common. Visiting a particular nest box could highlight a suitability check of the nesting area, and preference is shown if the nest box is already known. As we found, more visits are paid to nest boxes that were used by the prospecting female in the previous years. This showed an inclination for known and probably suitable territories. This did not mean that in the end the prospecting female succeeded in re-nesting (laying eggs for a second time in the season), and moreover doing it in the preferred nest site. The propensity of re-nesting after the first time in the season greatly depends on when the first clutch was started, as shown in the literature [37, 70] and in this study. But we were surprised to see an important contribution of the prospecting behaviour to predict the probability of having a second clutch. Furthermore, this links the prospecting behaviour to the annual fecundity of the females, in a way that more prospecting predicted second clutches and consequently more eggs laid per year. But also to the productivity, that for prospecting females did not increase despite having laid more eggs. In another study on jackdaws (*Corvus monedula*) higher prospecting individuals were having less fledglings than non-prospecting conspecifics [39]. The strategy adopted by prospecting females in our study does not seem to have a positive or negative effect on annual fledgling production, probably underlying neither positive nor negative selection, hence maintaining both prospecting and non-prospecting strategies in the population with a certain variation. The neutral outcome in productivity could be explained by the fact that raising a chick is a duty mostly carried out by male barn owls, so the female contribution to the fledgling final condition is secondary. In conclusion, we identified and associated prospecting behaviour to proximal causes and consequences in a nocturnal cosmopolitan species with the use of high-resolution biologging data. We high-lighted (1) the importance of such technology to identify and quantify behaviours remotely at night (i.e., prospecting), pairing landscape features (i.e., other nest boxes) with tracking data, (2) we linked the emergence and variation of the identified behaviour with individual- and partner-related variables including movement-related parameters (e.g., home range, activity budget), (3) we related the prospecting behaviour with individual fitness consequences that point at sex-related differential breeding strategies in a wild barn owl population.

## Acknowledgements

We would like to thank E. Millet, L. Ancay, A.P. Machado, R. Buhler, C. Gémard, C. Massa, A. C.Heinz, S. Zurkinden, N. Külling, M. Froehli and all the field assistants, students and interns from University of Lausanne and the Swiss Ornithological Institute for their help in collecting field data. This study was financially supported by the Swiss National Science Foundation (grants no. 310030_200321). We also want to thank all the members of the Barn Owl Research Group for feedback on this study during the last year, and P. Gibson for proofreading and editing.

## Authors contribution

Paolo Becciu conceived the study, analysed the data, and led the writing of the manuscript. Robin Séchaud, Kim Schalcher and Alexandre Roulin helped conceiving the study at early stage and collected the data. Robin Séchaud helped in data curation and calculated several variables used in the analyses. Céline Plancherel inspired this work with her MSc thesis project and gave fundamental comments improving the text. Alexandre Roulin funded the study and provided feedback on the interpretation of results. All authors contributed critically to the drafts and gave final approval for publication.

## References

[1] D. D. Brown, R. Kays, M. Wikelski, R. Wilson, and A. P. Klimley, “Observing the unwatchable through acceleration logging of animal behavior,” Animal Biotelemetry, vol. 1, no. 1, p. 1–16, 2013.

[2] R. Kays, M. C. Crofoot, W. Jetz, and M. Wikelski, “Terrestrial animal tracking as an eye on life and planet,” Science, vol. 348, no. 6240, p. aaa2478, 2015.

[3] R. Costa-Pereira, R. J. Moll, B. R. Jesmer, and W. Jetz, “Animal tracking moves community ecology: Opportunities and challenges,” Journal of Animal Ecology, 2022.

[4] R. Nathan, W. M. Getz, E. Revilla, M. Holyoak, R. Kadmon, D. Saltz, and P. E. Smouse, “A movement ecology paradigm for unifying organismal movement research,” Proceedings of the National Academy of Sciences of the United States of America, vol. 105, no. 49, p. 19052–19059, 2008.

[5] R. Nathan, C. T. Monk, R. Arlinghaus, T. Adam, J. Alós, M. As-saf, H. Baktoft, C. E. Beardsworth, M. G. Bertram, A. I. Bi-jleveld, T. Brodin, J. L. Brooks, A. Campos-Candela, S. J. Cooke, K. Gjelland, P. R. Gupte, R. Harel, G. Hellström, F. Jeltsch, S. S. Killen, T. Klefoth, R. Langrock, R. J. Lennox, E. Lourie, J. R. Madden, Y. Orchan, I. S. Pauwels, M. Říha, M. Roeleke, U. E. Schlägel, D. Shohami, J. Signer, S. Toledo, O. Vilk, S. Westrelin, M. A. Whiteside, and I. Jarić, “Big-data approaches lead to an increased understanding of the ecology of animal movement,” Science, vol. 375, Feb 2022.

[6] M. d. M. Delgado, V. Penteriani, E. Revilla, and V. O. Nams, “The effect of phenotypic traits and external cues on natal dispersal movements,” Journal of Animal Ecology, vol. 79, p. 620–632, May 2010.

[7] B. Doligez and T. Pärt, “Estimating fitness consequences of dispersal: a road to ‘know-where’? non-random dispersal and the underestimation of dispersers’ fitness,” Journal of Animal Ecology, vol. 77, p. 1199–1211, Nov 2008.

[8] R. E. Dunn, S. Wanless, F. Daunt, M. P. Harris, and J. A. Green, “A year in the life of a north atlantic seabird: behavioural and energetic adjustments during the annual cycle,” Scientific Reports, vol. 10, no. 5993, p. 1–11, 2020.

[9] R. Séchaud, K. Schalcher, B. Almasi, R. Bühler, K. Safi, A. Romano, and A. Roulin, “Home range size and habitat quality affect breeding success but not parental investment in barn owl males,” Scientific Reports, vol. 12, p. 6516, Dec 2022.

[10] M. Chimienti, F. M. Beest, L. T. Beumer, J. Desforges, L. H. Hansen, M. Stelvig, and N. M. Schmidt, “Quantifying behavior and life-history events of an arctic ungulate from year-long continuous accelerometer data,” Ecosphere, vol. 12, Jun 2021.

[11] L. M. San-Jose, R. Séchaud, K. Schalcher, C. Judes, A. Questiaux, A. Oliveira-Xavier, C. Gémard, B. Almasi, P. Béziers, A. Kelber, A. Amar, and A. Roulin, “Differential fitness effects of moonlight on plumage colour morphs in barn owls,” Nature Ecology and Evolution, vol. 3, no. 9, p. 1331–1340, 2019.

[12] M. Berlincourt, L. P. Angel, and J. P. Arnould, “Combined use of gps and accelerometry reveals fine scale three-dimensional foraging behaviour in the short-tailed shearwater,” PLoS ONE, vol. 10, no. 10, p. 1–16, 2015.

[13] C. Zavalaga, G. Dell’Omo, P. Becciu, and K. Yoda, “Patterns of gps tracks suggest nocturnal foraging by incubating peruvian pelicans (pelecanus thagus),” PLoS ONE, vol. 6, no. 5, 2011.

[14] S. Bar-David, I. Bar-David, P. C. Cross, S. J. Ryan, C. U. Knechtel, and W. M. Getz, “Methods for assessing movement path recursion with application to african buffalo in south africa,” Ecology, vol. 90, no. 9, 2009.

[15] N. Sapir, N. Horvitz, D. K. Dechmann, J. Fahr, and M. Wikelski, “Commuting fruit bats beneficially modulate their flight in relation to wind,” Proceedings of the Royal Society B: Biological Sciences, vol. 281, no. 1782, 2014.

[16] A. Tarroux, H. Weimerskirch, S. H. Wang, D. H. Bromwich, Y. Cherel, A. Kato, Y. Ropert-Coudert, Ø. Varpe, N. G. Yoccoz, and S. Descamps, “Flexible flight response to challenging wind conditions in a commuting antarctic seabird: Do you catch the drift?,” Animal Behaviour, vol. 113, no. January, p. 99–112, 2016.

[17] P. Becciu, S. Rotics, N. Horvitz, M. Kaatz, W. Fiedler, D. Zurell, A. Flack, F. Jeltsch, M. Wikelski, R. Nathan, and N. Sapir, “Causes and consequences of facultative sea crossing in a soaring migrant,” Functional Ecology, vol. 34, no. April, p. 840–852, 2020.

[18] H. J. Williams and K. Safi, “Certainty and integration of options in animal movement,” Trends in Ecology and Evolution, vol. 36, no. 11, pp. 990–999, 2021.

[19] C. Bracis, K. L. Bildstein, and T. Mueller, “Revisitation analysis uncovers spatio-temporal patterns in animal movement data,” Ecography, vol. 41, no. 11, p. 1801–1811, 2018.

[20] S. Picardi, B. J. Smith, M. E. Boone, P. C. Frederick, J. G. Cecere, D. Rubolini, L. Serra, S. Pirrello, R. R. Borkhataria, and M. Basille, “Analysis of movement recursions to detect reproductive events and estimate their fate in central place foragers,” Movement Ecology, vol. 8, no. 1, p. 1–14, 2020.

[21] U. E. Schlägel, J. Signer, A. Herde, S. Eden, F. Jeltsch, J. A. Eccard, and M. Dammhahn, “Estimating interactions between individuals from concurrent animal movements,” Methods in Ecology and Evolution, vol. 10, no. 8, p. 1234–1245, 2019.

[22] O. Berger-Tal and S. Bar-David, “Recursive movement patterns: Review and synthesis across species,” Ecosphere, vol. 6, no. 9, 2015.

[23] H. B. Brandl, S. C. Griffith, T. Laaksonen, and W. Schuett, “Begging calls provide social cues for prospecting conspecifics in the wild zebra finch (taeniopygia guttata),” Auk, vol. 136, Apr 2019.

[24] T. Dittmann and P. H. Becker, “Sex, age, experience and condition as factors affecting arrival date in prospecting common terns, sterna hirundo,” Animal Behaviour, vol. 65, no. 5, p. 981–986, 2003.

[25] D. Oro, J. Bécares, F. Bartumeus, and J. M. Arcos, “High frequency of prospecting for informed dispersal and colonisation in a social species at large spatial scale,” Oecologia, vol. 197, p. 395–409, Oct 2021.

[26] T. Pärt, D. Arlt, B. Doligez, M. Low, and A. Qvarnström, “Prospectors combine social and environmental information to improve habitat selection and breeding success in the subsequent year,” Journal of Animal Ecology, vol. 80, p. 1227–1235, Nov 2011.

[27] J.-F. Therrien, D. Pinaud, G. Gauthier, N. Lecomte, K. L. Bildstein, and J. Bety, “Is pre-breeding prospecting behaviour affected by snow cover in the irruptive snowy owl? a test using statespace modelling and environmental data annotated via movebank,” Movement Ecology, vol. 3, p. 1, Dec 2015.

[28] T. Boulinier and E. Danchin, “The use of conspecific reproductive success for breeding patch selection in terrestrial migratory species,” Evolutionary Ecology, vol. 11, 1997.

[29] L. Giraldeau, T. J. Valone, and J. J. Templeton, “Potential disadvantages of using socially acquired information,” Philosophical Transactions of the Royal Society of London. Series B: Biological Sciences, vol. 357, p. 1559–1566, Nov 2002.

[30] T. Boulinier, K. D. McCoy, N. G. Yoccoz, J. Gasparini, and T. Tveraa, “Public information affects breeding dispersal in a colonial bird: Kittiwakes cue on neighbours,” Biology Letters, vol. 4, p. 538–540, Oct 2008.

[31] M. Ciaglo, R. Calhoun, S. W. Yanco, M. B. Wunder, C. A. Stricker, and B. D. Linkhart, “Evidence of postbreeding prospecting in a long-distance migrant,” Ecology and Evolution, vol. 11, no. 1, p. 599–611, 2021.

[32] M. C. Zicus and S. K. Hennes, “Nest prospecting by common goldeneyes,” The Condor, vol. 91, no. 4, 1989.

[33] A. Ponchon, T. Chambert, E. Lobato, T. Tveraa, D. Grémillet, and T. Boulinier, “Breeding failure induces large scale prospecting movements in the black-legged kittiwake,” Journal of Experimental Marine Biology and Ecology, vol. 473, p. 138–145, Dec 2015.

[34] A. Ponchon, L. Iliszko, D. Grémillet, T. Tveraa, and T. Boulinier, “Intense prospecting movements of failed breeders nesting in an unsuccessful breeding subcolony,” Animal Behaviour, vol. 124, p. 183–191, Feb 2017.

[35] D. Parejo, T. Pérez-Contreras, C. Navarro, J. J. Soler, and J. M. Avilés, “Spotless starlings rely on public information while visiting conspecific nests: an experiment,” Animal Behaviour, vol. 75, p. 483–488, Feb 2008.

[36] A. Roulin, Barn Owls. Cambridge: Cambridge University Press, 2020.

[37] P. Béziers and A. Roulin, “Double brooding and offspring desertion in the barn owl tyto alba,” Journal of Avian Biology, vol. 47, no. 2, p. 235–244, 2016.

[38] R. Séchaud, K. Schalcher, A. P. Machado, B. Almasi, C. Massa, K. Safi, and A. Roulin, “Behaviour-specific habitat selection patterns of breeding barn owls,” Movement Ecology, vol. 9, no. 1, p. 1–11, 2021.

[39] W. Schuett, J. Laaksonen, and T. Laaksonen, “Prospecting at conspecific nests and exploration in a novel environment are associated with reproductive success in the jackdaw,” Behavioral Ecology and Sociobiology, vol. 66, p. 1341–1350, Sep 2012.

[40] R. C. Fijn, P. Wolf, W. Courtens, H. Verstraete, E. W. Stienen, L. Iliszko, and M. J. Poot, “Post-breeding prospecting trips of adult sandwich terns thalasseus sandvicensis,” Bird Study, vol. 61, p. 566–571, Oct 2014.

[41] C. Frey, C. Sonnay, A. Dreiss, and A. Roulin, “Habitat, breeding performance, diet and individual age in swiss barn owls (tyto alba),” Journal of Ornithology, vol. 152, p. 279–290, Apr 2011.

[42] M. K. Labocha and J. P. Hayes, “Morphometric indices of body condition in birds: A review,” Journal of Ornithology, vol. 153, no. 1, p. 1–22, 2012.

[43] A. L. Ducrest, L. Keller, and A. Roulin, “Pleiotropy in the melanocortin system, coloration and behavioural syndromes,” Trends in Ecology and Evolution, vol. 23, p. 502–510, Sep 2008.

[44] A. Roulin and R. Altwegg, “Breeding rate is associated with pheomelanism in male and with eumelanism in female barn owls,” Behavioral Ecology, vol. 18, no. 3, p. 563–570, 2007.

[45] A. Roulin and A. L. Ducrest, “Association between melanism, physiology and behaviour: A role for the melanocortin system,” European Journal of Pharmacology, vol. 660, p. 226–233, Jun 2011.

[46] P. Karell, K. Kohonen, and K. Koskenpato, “Specialist predation covaries with colour polymorphism in tawny owls,” Behavioral Ecology and Sociobiology, vol. 75, no. 3, 2021.

[47] A. Romano, R. Séchaud, A. H. Hirzel, and A. Roulin, “Climate-driven convergent evolution of plumage colour in a cosmopolitan bird,” Global Ecology and Biogeography, vol. 28, no. 4, p. 496–507, 2019.

[48] V. Amrhein and S. Greenland, “Rewriting results in the language of compatibility,” Trends in Ecology and Evolution, vol. 37, no. 7, pp. 567–568, 2022. Special issue: Symbiosis.

[49] S. Muff, E. B. Nilsen, R. B. O’Hara, and C. R. Nater, “Rewriting results sections in the language of evidence,” Trends in Ecology and Evolution, vol. 37, p. 203–210, Mar 2022.

[50] A. F. Zuur, E. N. Ieno, and C. S. Elphick, “A protocol for data exploration to avoid common statistical problems,” Methods in Ecology and Evolution, vol. 1, no. 1, p. 3–14, 2010.

[51] K. P. Anderson and D. A. Burnham, Model Selection and Multi-Model Inference: A Practical Information-Theoretic Approach (2nd Edition), vol. 172. Springer, 2002.

[52] N. Sugiura, “Further analysts of the data by akaike’s information criterion and the finite corrections,” Communications in Statistics - Theory and Methods, vol. 7, no. 1, p. 13–26, 1978.

[53] C. E. Grueber, S. Nakagawa, R. J. Laws, and I. G. Jamieson, “Multimodel inference in ecology and evolution: Challenges and solutions,” Journal of Evolutionary Biology, vol. 24, no. 4, p. 699–711, 2011.

[54] S. Nakagawa and R. P. Freckleton, “Model averaging, missing data and multiple imputation: a case study for behavioural,” Source: Behavioral Ecology and Sociobiology, vol. 65, no. 1, p. 103–116, 2011.

[55] D. R. Anderson, K. P. Burnham, and W. L. Thompson, “Null hypothesis testing: Problems, prevalence, and an alternative,” The Journal of Wildlife Management, vol. 64, no. 4, 2000.

[56] A. F. Zuur, E. N. Ieno, N. Walker, A. A. Saveliev, and G. M. Smith, Mixed effects models and extension in ecology with R. New York: Springer, 2009.

[57] F. Hartig, “Residual diagnostics for hierarchical models,” 2019.

[58] R. C. Team, “R: A language and environment for statistical computing,” no. 4.0.2, 2020.

[59] M. E. Brooks, K. Kristensen, K. J. van Benthem, A. Magnusson, C. W. Berg, A. Nielsen, H. J. Skaug, M. Mächler, and B. M. Bolker, “glmmtmb balances speed and flexibility among packages for zero-inflated generalized linear mixed modeling,” R Journal, vol. 9, no. 2, p. 378–400, 2017.

[60] K. Barton, “Mumin: Multi-model inference, version 1.43.6,” R Package Version 1..42..1, no. 1, p. 1–75, 2019.

[61] R. Team, “Rstudio: Integrated development for r,” 2020.

[62] M. del Mar Delgado, I. I. Ratikainen, and H. Kokko, “Inertia: the discrepancy between individual and common good in dispersal and prospecting behaviour,” Biological Reviews, vol. 86, p. 717–732, Aug 2011.

[63] J. M. Reed, T. Boulinier, E. Danchin, and L. W. Oring, Informed dispersal. Prospecting by Birds for Breeding Sites, vol. 15, ch. 5. New York: Kluwer Academic / Plenum Publishers, 1999.

[64] A. Roulin, C. Riols, C. Dijkstra, and A. L. Ducrest, “Female plumage spottiness signals parasite resistance in the barn owl (tyto alba),” Behavioral Ecology, vol. 12, no. 1, p. 103–110, 2001.

[65] O. Peleg, M. Charter, Y. Leshem, I. Izhaki, and A. Roulin, “Conditional association between melanism and personality in israeli barn owls,” Bird Study, vol. 61, p. 572–577, Oct 2014.

[66] A. Roulin, “Nonrandom pairing by male barn owls (tyto alba) with respect to a female plumage trait,” Behavioral Ecology, vol. 10, no. 6, p. 688–695, 1999.

[67] B. Almasi, C. Massa, L. Jenni, and A. Roulin, “Exogenous corticosterone and melanin-based coloration explain variation in juvenile dispersal behaviour in the barn owl (*Tyto alba*),” PLoS ONE, vol. 16, p. e0256038, Sep 2021.

[68] K. A. Leighty, J. Soltis, C. M. Wesolek, and A. Savage, “Rumble vocalizations mediate interpartner distance in african elephants, loxodonta africana,” Animal Behaviour, vol. 76, p. 1601–1608, Nov 2008.

[69] M. J. Noonan, R. Martinez-Garcia, G. H. Davis, M. C. Crofoot, R. Kays, B. T. Hirsch, D. Caillaud, E. Payne, A. Sih, D. L. Sinn, O. Spiegel, W. F. Fagan, C. H. Fleming, and J. M. Calabrese, “Estimating encounter location distributions from animal tracking data,” Methods in Ecology and Evolution, vol. 12, p. 1158–1173, Jul 2021.

[70] A. Roulin, “Offspring desertion by double-brooded female barn owls (*Tyto Alba*),” The Auk, vol. 119, no. 2, 2002.

