## Supplementary material for "Movement data link phenotypic traits to individual fitness in a nocturnal predator": Fig. S1, Fig. S2, Fig. S3

### Electronic Supplementary Materials

#### GPS tags, deployment, behaviour annotation and home range size

Barn owls were tagged using GiPSy-5 GPS tags (2016-2017) and Axy-Trek GPS and accelerometer tags (2018-2020) (https://www.technosmart.eu/, Technosmart, Italy) weighing approximately 12 g and 12.5 g including battery, respectively (less than 5% of the owl body mass; in our population, body mass ranged from 251 to 393 g). They were attached as backpacks with a Teflon harness. Each tag collected location and time every 10 s (GiPSy-5) or every second (Axy-Trek), from 30 min before dusk until 30 min after dawn, covering the entire owl nocturnal activity period. Axy-Trek tracks were resampled at 10 s with the function *thin_track* by the package "move" (Kranstauber et al., 2018) in the R environment (R Core Team, 2020). Breeding barn owls were captured at their nest site when the oldest offspring was 25 days-old on average (SD = 2.8), equipped with GPS tags and released at the capture site. We recorded adults' sex and age (categorized as yearlings or older birds). Approximately two weeks later, the owls were recaptured and the GPS tags recovered. The deployment of GPS tags corresponds to a period of intense hunting effort by the parents to feed their nestlings, while being in accordance with ethical (earlier captures could lead to the abandonment of the clutch) and methodological constraints (later recapture could be compromised due to changes in food provisioning behaviour). Prior to any analysis, the tracks were filtered for aberrant positions using speed (excluding locations with a speed higher than 15 m/s) and location (excluding locations outside the study area). Tagged individuals were weighed at the moment of their capture, both times when the data-logger was deployed and retrieved. Also, chicks were counted and weighed, and measurement of wing length were taken.

We annotated the tracks using the Expectation-Maximization binary Clustering (EMbC) method implemented in the "EMbC" package (Garriga et al., 2016). The three behavioural modes distinguished were perching, hunting, and commuting. Perching, as a stationary behaviour, was characterized by low speed and a wide range of turning angles due to small GPS location errors. Hunting was characterized by low-medium speed and medium-high turning angles, whereas commuting was characterised by fast and straight flights. For validation, the EMbC behavioural classification was confronted to a visual classification performed on the tracks of 20 individuals. We found an average match of 92.7% between the visual and the EMbC classifications (perching: mean = 94.5%, SE = 2.3; hunting: mean = 92.6%, SE = 4.9; commuting: mean = 91.1%, SE = 3.8) (San-Jose et al., 2019; Séchaud et al., 2021).

Home range size was calculated using a 95% kernel density estimator method (Worton, 1989). To deal with temporal autocorrelation between data points, we used the continuous-time movement modelling package "ctmm" (Calabrese et al., 2016) to calculate home range size via auto-correlated kernel density estimation (AKDE) (Fleming et al., 2015). The ctmm model was calibrated using User Equivalent Range Error (UERE), estimated with location data obtained by fixed GPS devices in an open landscape. Model parameters with better fit were chosen automatically with the function *variogram.fit* in the "ctmm" package (Calabrese et al., 2016).

#### References


Calabrese, J. M., Fleming, C. H., & Gurarie, E. (2016). Ctmm: an R Package for Analyzing Animal Relocation Data As a Continuous-Time Stochastic Process. *Methods in Ecology and Evolution*, *7*(9), 1124–1132. https://doi.org/10.1111/2041-210X.12559

Fleming, C. H., Fagan, W. F., Mueller, T., Olson, K. A., Leimgruber, P., & Calabrese, J. M. (2015). Rigorous home range estimation with movement data: a new autocorrelated kernel density estimator. *Ecology*, *96*(5), 1182–1188. https://doi.org/10.1890/14-2010.1

Garriga, J., Palmer, J. R. B., Oltra, A., & Bartumeus, F. (2016). Expectation-maximization binary clustering for behavioural annotation. *PLoS ONE*, *11*(3), 1–26. https://doi.org/10.1371/journal.pone.0151984

Kranstauber, B., Smolla, M., & Scharf, A. K. (2018). move: Visualizing and Analyzing Animal Track Data Version. *R Package Version 3.1.0.* https://cran.r-project.org/package=move

R Core Team. (2020). *R: A language and environment for statistical computing* (4.0.2). R Foundation for Statistical Computing. https://www.r-project.org/

San-Jose, L. M., Séchaud, R., Schalcher, K., Judes, C., Questiaux, A., Oliveira-Xavier, A., Gémard, C., Almasi, B., Béziers, P., Kelber, A., Amar, A., & Roulin, A. (2019). Differential fitness effects of moonlight on plumage colour morphs in barn owls. *Nature Ecology and Evolution*, *3*(9), 1331–1340. https://doi.org/10.1038/s41559-019-0967-2

Séchaud, R., Schalcher, K., Machado, A. P., Almasi, B., Massa, C., Safi, K., & Roulin, A. (2021). Behaviour-specific habitat selection patterns of breeding barn owls. *Movement Ecology*, *9*(1), 1–11. https://doi.org/10.1186/s40462-021-00258-6

Worton, B. J. (1989). Kernel Methods for Estimating the Utilization Distribution in Home-Range Studies. *Ecology*, *70*(1), 164–168. https://doi.org/10.2307/1938423

Correlation between diameter and number (per unit of surface – 60x40 mm frame) of spots on breast and belly

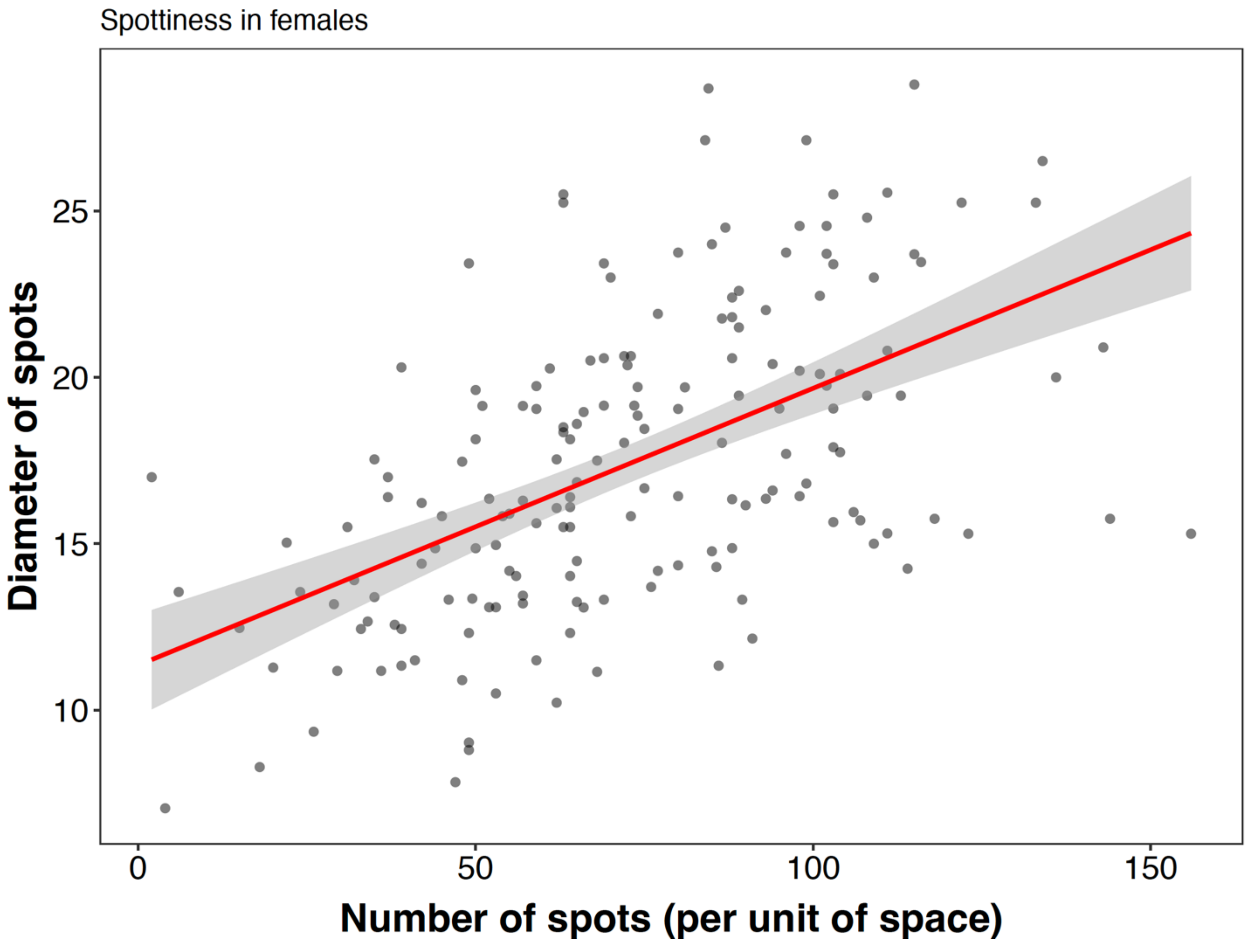


**Figure S1** – Correlation between number of melanic spots and their diameter. Pearson's correlation: $r = 0.54$, $t = 8.43$, $df = 171$, $P < 0.0001$, $n = 173$.

Distribution of tagging duration among individuals

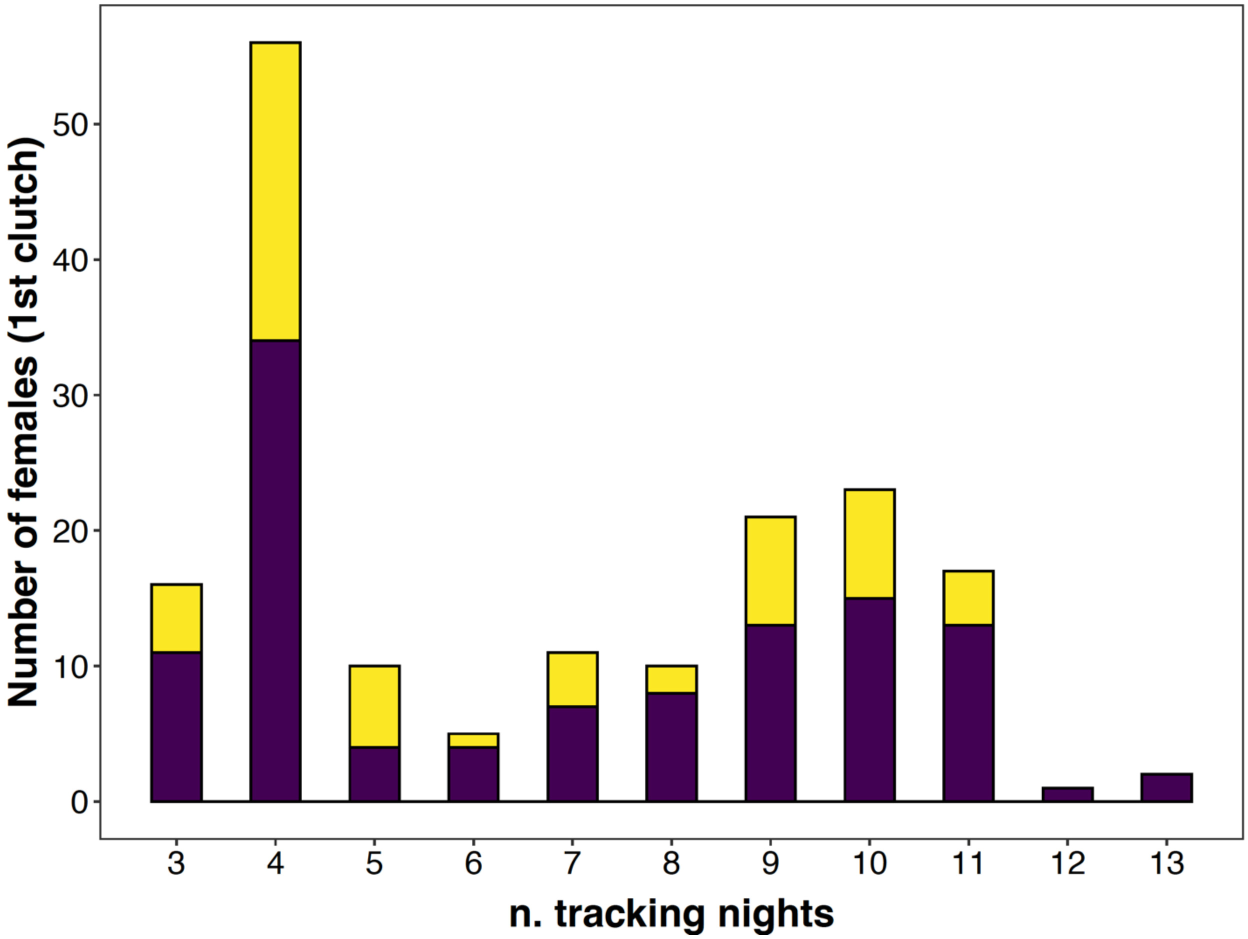


**Figure S2** – Distribution of tracking nights among tagged female barn owls considered in the analyses. In yellow are the females that do not visit any other nest box during the tracking period. In violet are the females that visit at least one nest box.

#### Averaged-model coefficients and 95% C.I. related to Figure 2 in main text

Brood-related predictors (n = 116 )

| ***response****/predictors* | *coef* | *lowerCI* | *upperCI* | *sum.weights* |
|---|---|---|---|---|
| **prospecting[0,1]** (binomial distr.) | | | | |
| Brood size | -0.20 | -0.63 | 0.23 | 0.35 |
| Wing growth rate | -0.22 | -1.10 | 0.66 | 0.28 |
| Brood size diff. tagging vs recovery | 0.09 | -0.40 | 0.59 | 0.28 |
| Laying date | -0.03 | -0.43 | 0.37 | 0.26 |

* Null hypothesis value outside the confidence interval.

**Table S1** – Summary table of averaged coefficients and 95% confidence intervals and variable importance (sum of Aikike weights) for the brood-related effects on binomial variable of prospecting (1 = female barn owl prospected at least one other nest box, 0 = no visits to other nest boxes).

Female-related predictors (n = 135 )

| ***response****/predictors* | *coef* | *lowerCI* | *upperCI* | *sum.weights* |
|---|---|---|---|---|
| **prospecting[0,1]** (binomial distr.) | | | | |
| Spottiness | -0.54* | -0.93 | -0.14 | 0.95 |
| Age | 0.53 | -0.24 | 1.31 | 0.47 |
| Daily mass variation | 0.26 | -0.15 | 0.67 | 0.45 |
| Body condition (resid.) | -0.13 | -0.54 | 0.28 | 0.31 |
| Colour | 0.07 | -0.34 | 0.47 | 0.27 |

* Null hypothesis value outside the confidence interval.

**Table S2** – Summary table of averaged coefficients and 95% confidence intervals and variable importance (sum of Aikike weights) for the female-related effects on binomial variable of prospecting (1 = female barn owl prospected at least one other nest box, 0 = no visits to other nest boxes).

Male-related predictors (n = 122 )

| ***response****/predictors* | *coef* | *lowerCI* | *upperCI* | *sum.weights* |
|---|---|---|---|---|
| **prospecting[0,1]** (binomial distr.) | | | | |

| | | | | |
|---|---|---|---|---|
| Spottiness | -0.29 | -0.69 | 0.11 | 0.50 |
| Provisioning rate | 0.19 | -0.23 | 0.60 | 0.34 |
| Age | -0.33 | -1.12 | 0.47 | 0.32 |
| Body condition (resid.) | 0.16 | -0.24 | 0.56 | 0.32 |
| Time spent perching | -0.12 | -0.52 | 0.29 | 0.29 |
| Home range size | -0.10 | -0.53 | 0.32 | 0.28 |
| Colour | -0.07 | -0.52 | 0.38 | 0.28 |

* Null hypothesis value outside the confidence interval.

**Table S3** – Summary table of averaged coefficients and 95% confidence intervals and variable importance (sum of Aikike weights) for the male-related effects on binomial variable of prospecting (1 = female barn owl prospected at least one other nest box, 0 = no visits to other nest boxes).

Brood-related predictors (n = 116)

| ***response****/predictors* | *coef* | *lowerCI* | *upperCI* | *sum.weights* |
|---|---|---|---|---|
| **Mean visits** (tweedie distr.) | | | | |
| Laying date | -0.21 | -0.45 | 0.03 | 0.59 |
| Brood size | -0.17 | -0.39 | 0.05 | 0.52 |
| Wing growth rate | -0.29 | -0.73 | 0.15 | 0.44 |
| Brood size diff. tagging vs recovery | -0.03 | -0.32 | 0.26 | 0.28 |
| **n. nest sites visited** (Negative binomial distr.) | | | | |
| Brood size | -0.17 | -0.39 | 0.04 | 0.56 |
| Laying date | -0.10 | -0.33 | 0.13 | 0.34 |
| Wing growth rate | -0.14 | -0.57 | 0.29 | 0.30 |
| Brood size diff. tagging vs recovery | -0.02 | -0.29 | 0.26 | 0.28 |
| **median time at nest sites** (tweedie distr.) | | | | |
| Wing growth rate | 0.47* | 0.01 | 0.94 | 0.72 |
| Brood size diff. tagging vs recovery | 0.28 | -0.01 | 0.57 | 0.70 |
| Laying date | -0.23 | -0.48 | 0.02 | 0.64 |

| | | | | |
|---|---|---|---|---|
| Brood size | -0.24 | -0.51 | 0.03 | 0.63 |

* Null hypothesis value outside the confidence interval.

**Table S4** – Summary table of averaged coefficients and 95% confidence intervals and variable importance (sum of Aikike weights) for the brood-related effects on the three response variables considered (prospecting parameters).

Female-related predictors (n = 135)

| ***response****/predictors* | *coef* | *lowerCI* | *upperCI* | *sum.weights* |
|---|---|---|---|---|
| **Mean visits** (tweedie distr.) | | | | |
| Spottiness | -0.28* | -0.48 | -0.08 | 0.93 |
| Age | 0.32 | -0.12 | 0.76 | 0.49 |
| Colour | -0.05 | -0.26 | 0.16 | 0.28 |
| Body condition (resid.) | -0.03 | -0.26 | 0.19 | 0.26 |
| Daily mass variation | -0.02 | -0.24 | 0.21 | 0.26 |
| **n. nest sites visited** (Negative binomial distr.) | | | | |
| Spottiness | -0.29* | -0.49 | -0.09 | 0.97 |
| Colour | 0.14 | -0.06 | 0.33 | 0.48 |
| Age | 0.27 | -0.14 | 0.68 | 0.45 |
| Body condition (resid.) | -0.04 | -0.24 | 0.17 | 0.27 |
| Daily mass variation | 0.01 | -0.19 | 0.21 | 0.26 |
| **median time at nest sites** (tweedie distr.) | | | | |
| Spottiness | -0.41* | -0.62 | -0.20 | 1.00 |
| Daily mass variation | 0.22 | -0.00 | 0.44 | 0.70 |
| Body condition (resid.) | 0.15 | -0.08 | 0.39 | 0.45 |
| Colour | 0.07 | -0.12 | 0.27 | 0.31 |
| Age | -0.15 | -0.57 | 0.28 | 0.30 |

* Null hypothesis value outside the confidence interval.

**Table S5** – Summary table of averaged coefficients and 95% confidence intervals and variable importance (sum of Aikike weights) for the female-related effects on the three response variables considered (prospecting parameters).

Male-related predictors (n = 122)

| ***response**/predictors* | *coef* | *lowerCI* | *upperCI* | *sum.weights* |
|---|---|---|---|---|
| **Mean visits** (tweedie distr.) | | | | |
| Home range size | -0.32* | -0.52 | -0.11 | 0.97 |
| Body condition (resid.) | 0.20* | 0.02 | 0.38 | 0.76 |
| Provisioning rate | 0.19 | -0.00 | 0.38 | 0.68 |
| Spottiness | -0.17 | -0.39 | 0.05 | 0.53 |
| Age | -0.24 | -0.65 | 0.16 | 0.39 |
| Colour | 0.07 | -0.18 | 0.31 | 0.29 |
| Time spent perching | -0.03 | -0.27 | 0.21 | 0.26 |
| **n. nest sites visited** (Negative binomial distr.) | | | | |
| Home range size | -0.26* | -0.47 | -0.05 | 0.88 |
| Spottiness | -0.17 | -0.37 | 0.03 | 0.57 |
| Time spent perching | -0.16 | -0.37 | 0.05 | 0.52 |
| Provisioning rate | 0.13 | -0.07 | 0.32 | 0.43 |
| Body condition (resid.) | 0.09 | -0.09 | 0.27 | 0.36 |
| Age | -0.17 | -0.55 | 0.22 | 0.32 |
| Colour | 0.01 | -0.22 | 0.24 | 0.26 |
| **median time at nest sites** (tweedie distr.) | | | | |
| Provisioning rate | 0.31* | 0.13 | 0.48 | 0.99 |
| Body condition (resid.) | -0.30* | -0.50 | -0.11 | 0.98 |
| Home range size | 0.11 | -0.12 | 0.35 | 0.35 |
| Spottiness | -0.10 | -0.31 | 0.11 | 0.34 |
| Time spent perching | -0.10 | -0.32 | 0.12 | 0.33 |
| Colour | 0.04 | -0.20 | 0.27 | 0.26 |
| Age | 0.04 | -0.38 | 0.45 | 0.25 |

* Null hypothesis value outside the confidence interval.

**Table S6** – Summary table of averaged coefficients and 95% confidence intervals and variable importance (sum of Aikike weights) for the male-related effects on the three response variables considered (prospecting parameters).

#### Models considering only females that visit at least one nest box

Brood-related predictors (n = 76)

| ***response***/*predictors* | *coef* | *lowerCI* | *upperCI* | *sum.weights* |
|---|---|---|---|---|
| **Mean visits** (log transf. – gaussian distr.) | | | | |
| Laying date | -0.19 | -0.38 | 0.01 | 0.64 |
| Brood size | -0.15 | -0.35 | 0.06 | 0.48 |
| Wing growth rate | -0.11 | -0.29 | 0.08 | 0.39 |
| Brood size diff. tagging vs recovery | -0.06 | -0.32 | 0.19 | 0.30 |
| **n. nest sites visited** (log transf. – gaussian distr.) | | | | |
| Brood size | -0.10 | -0.26 | 0.05 | 0.44 |
| Laying date | -0.06 | -0.21 | 0.09 | 0.31 |
| Brood size diff. tagging vs recovery | -0.01 | -0.19 | 0.18 | 0.27 |
| Wing growth rate | 0.01 | -0.13 | 0.15 | 0.25 |
| **median time at nest sites** (log transf. – gaussian distr.) | | | | |
| Laying date | -0.21* | -0.39 | -0.02 | 0.78 |
| Wing growth rate | 0.12 | -0.05 | 0.29 | 0.46 |
| Brood size | -0.11 | -0.32 | 0.11 | 0.35 |
| Brood size diff. tagging vs recovery | 0.09 | -0.16 | 0.34 | 0.31 |

* Null hypothesis value outside the confidence interval.

**Table S7** – Summary table of averaged coefficients and 95% confidence intervals and variable importance (sum of Aikike weights) for the brood-related effects on the three response variables considered (prospecting parameters) but without the females that do not visit any other nest box.

Female-related predictors (n = 88)

| ***response***/*predictors* | *coef* | *lowerCI* | *upperCI* | *sum.weights* |
|---|---|---|---|---|
| **Mean visits** (log transf. – gaussian distr.) | | | | |
| Spottiness | -0.10 | -0.29 | 0.08 | 0.39 |
| Daily mass variation | -0.07 | -0.26 | 0.12 | 0.31 |

| | | | | |
|---|---|---|---|---|
| Body condition (resid.) | 0.05 | -0.14 | 0.23 | 0.27 |
| Colour | -0.01 | -0.19 | 0.17 | 0.25 |
| Age | 0.03 | -0.36 | 0.41 | 0.25 |
| **n. nest sites visited** (log transf. – gaussian distr.) | | | | |
| Spottiness | -0.09 | -0.23 | 0.04 | 0.46 |
| Colour | 0.09 | -0.05 | 0.22 | 0.43 |
| Daily mass variation | -0.06 | -0.20 | 0.07 | 0.33 |
| Body condition (resid.) | 0.04 | -0.10 | 0.17 | 0.28 |
| Age | 0.02 | -0.27 | 0.30 | 0.25 |
| **median time at nest sites** (log transf. – gaussian distr.) | | | | |
| Spottiness | -0.16 | -0.33 | 0.01 | 0.65 |
| Age | -0.28 | -0.63 | 0.08 | 0.52 |
| Body condition (resid.) | 0.08 | -0.10 | 0.26 | 0.34 |
| Colour | -0.04 | -0.23 | 0.14 | 0.28 |
| Daily mass variation | 0.05 | -0.13 | 0.23 | 0.28 |

* Null hypothesis value outside the confidence interval.

**Table S8** – Summary table of averaged coefficients and 95% confidence intervals and variable importance (sum of Aikike weights) for the female-related effects on the three response variables considered (prospecting parameters) but without the females that do not visit any other nest box.

**Male-related predictors (n = 81)**

| ***response***/*predictors* | *coef* | *lowerCI* | *upperCI* | *sum.weights* |
|---|---|---|---|---|
| **Mean visits** (log transf. – gaussian distr.) | | | | |
| Provisioning rate | 0.20* | 0.02 | 0.39 | 0.77 |
| Body condition (resid.) | 0.18* | 0.01 | 0.35 | 0.72 |
| Home range size | -0.19 | -0.37 | -0.00 | 0.70 |
| Spottiness | -0.11 | -0.29 | 0.07 | 0.41 |
| Age | -0.18 | -0.53 | 0.17 | 0.35 |
| Time spent perching | 0.05 | -0.16 | 0.26 | 0.28 |

| | | | | |
|---|---|---|---|---|
| Colour | 0.03 | -0.16 | 0.22 | 0.25 |
| **n. nest sites visited** (log transf. – gaussian distr.) | | | | |
| Home range size | -0.17* | -0.31 | -0.02 | 0.82 |
| Time spent perching | -0.10 | -0.25 | 0.05 | 0.46 |
| Provisioning rate | 0.08 | -0.06 | 0.22 | 0.37 |
| Age | -0.14 | -0.40 | 0.12 | 0.36 |
| Body condition (resid.) | 0.07 | -0.06 | 0.19 | 0.35 |
| Spottiness | -0.06 | -0.19 | 0.07 | 0.32 |
| Colour | 0.00 | -0.14 | 0.14 | 0.24 |
| **median time at nest sites** (log transf. – gaussian distr.) | | | | |
| Provisioning rate | 0.25* | 0.09 | 0.42 | 0.96 |
| Body condition (resid.) | -0.24* | -0.40 | -0.08 | 0.95 |
| Home range size | 0.12 | -0.05 | 0.29 | 0.45 |
| Colour | 0.09 | -0.08 | 0.26 | 0.36 |
| Time spent perching | -0.01 | -0.20 | 0.17 | 0.25 |
| Age | 0.06 | -0.27 | 0.39 | 0.25 |
| Spottiness | 0.01 | -0.16 | 0.18 | 0.24 |

* Null hypothesis value outside the confidence interval.

**Table S9** – Summary table of averaged coefficients and 95% confidence intervals and variable importance (sum of Aikike weights) for the male-related effects on the three response variables considered (prospecting parameters) but without the females that do not visit any other nest box.

#### Binomial GLM summary relative to Fig. 3 in the main text

**Probability of multiple visits to a nest box** (1 = 1+ visits, 0 = only one visit)

| *Predictors* | *Odds Ratios* | *Conf.Int. (2.5%-97.5%)* |
|---|---|---|
| nest active previous years | 1.076 | 0.767 – 1.510 |
| mean fledglings previous years | 1.170 | 0.821 – 1.668 |
| breeding attempts previous years [1] | 4.763*** | 2.288 – 9.913 |

| | | |
|---|---|---|
| nest active current year [1] | 1.032 | 0.512 – 2.077 |
| nest distance (log) | 0.665* | 0.480 – 0.919 |
| nest density | 0.954 | 0.679 – 1.339 |
| Observations | | 233 |

* Null hypothesis value outside the confidence interval.
** p<0.05 ** p<0.01 *** p<0.001*

**Table S10** – Summary table of GLM showing Odds Ratios and 95% C.I. of predictors effect on probability of having multiple visit to a specific nest box [1] or only one [0] (binomial variable).

#### Summary of univariate binomial logistic regression models represented in Fig. 4A

| Prob. of re-nesting | | | | |
|---|---|---|---|---|
| *Predictors* | *Odds Ratios* | *Conf.Int. (2.5%-97.5%)* | *AICc* | $R^2$ |
| prospecting[0,1] [1] | 1.768 | 0.794 – 3.936 | 185.65 | 0.022 |
| Mean visits | 1.531* | 1.105 – 2.121 | 181.18 | 0.052 |
| n. nest sites visited | 1.435* | 1.031 – 1.998 | 183.21 | 0.038 |
| median time at nest sites | 1.346 | 0.978 – 1.853 | 184.50 | 0.026 |
| Observations | | | | 171 |

* Null hypothesis value outside the confidence interval.
** p<0.05 ** p<0.01 *** p<0.001*

**Table S11** – Summary table of four univariate models having the binomial variable re-nesting [1] or not [0] as response variable. This table highlights the predictive difference between predictors through AICc, showing Odds Ratios and 95% C.I..

#### Summary of averaged model represented in Fig. 4B

| probability of re-nesting (n = 127) | | | | |
|---|---|---|---|---|
| *Predictors* | *coef* | *lowerCI* | *upperCI* | *sum.weights* |
| Mean visits | 0.42* | 0.06 | 0.78 | 0.82 |
| Age (F) | -0.47 | -1.38 | 0.44 | 0.37 |
| Brood size | -0.13 | -0.57 | 0.31 | 0.30 |

| | | | | |
|---|---|---|---|---|
| Home range size (M) | -0.08 | -0.53 | 0.37 | 0.27 |
| Fledglings to nestlings ratio | -0.05 | -0.54 | 0.44 | 0.26 |
| AICc = 139.77 / $R^2$ = 0.08 | | | | |

* Null hypothesis value outside the confidence interval

**Table S12** – Summary table of averaged coefficients and 95% confidence intervals and variable importance (sum of Aikike weights) for the selected predictors on the re-nesting binomial response variable considered (re-nesting [1] or not [0]).

#### Summary of the averaged model predicting "probability of re-nesting" including "laying date" as predictor

| probability of re-nesting (n = 127) | | | | |
|---|---|---|---|---|
| *Predictors* | *coef* | *lowerCI* | *upperCI* | *sum.weights* |
| Laying date | -1.74* | -2.50 | -0.97 | 1.00 |
| Brood size | -0.45 | -1.02 | 0.11 | 0.57 |
| Mean visits | 0.33 | -0.12 | 0.77 | 0.51 |
| Age (F) | -0.42 | -1.46 | 0.63 | 0.32 |
| Home range size (M) | 0.14 | -0.39 | 0.67 | 0.29 |
| Fledglings to nestlings ratio | 0.04 | -0.58 | 0.66 | 0.27 |
| AICc = 110.51 / $R^2$ = 0.51 | | | | |

* Null hypothesis value outside the confidence interval

**Table S13** – Summary table of averaged coefficients and 95% confidence intervals and variable importance (sum of Aikike weights) for the selected predictors on the re-nesting binomial response variable considered (re-nesting [1] or not [0]). This model includes the laying date as predictor.

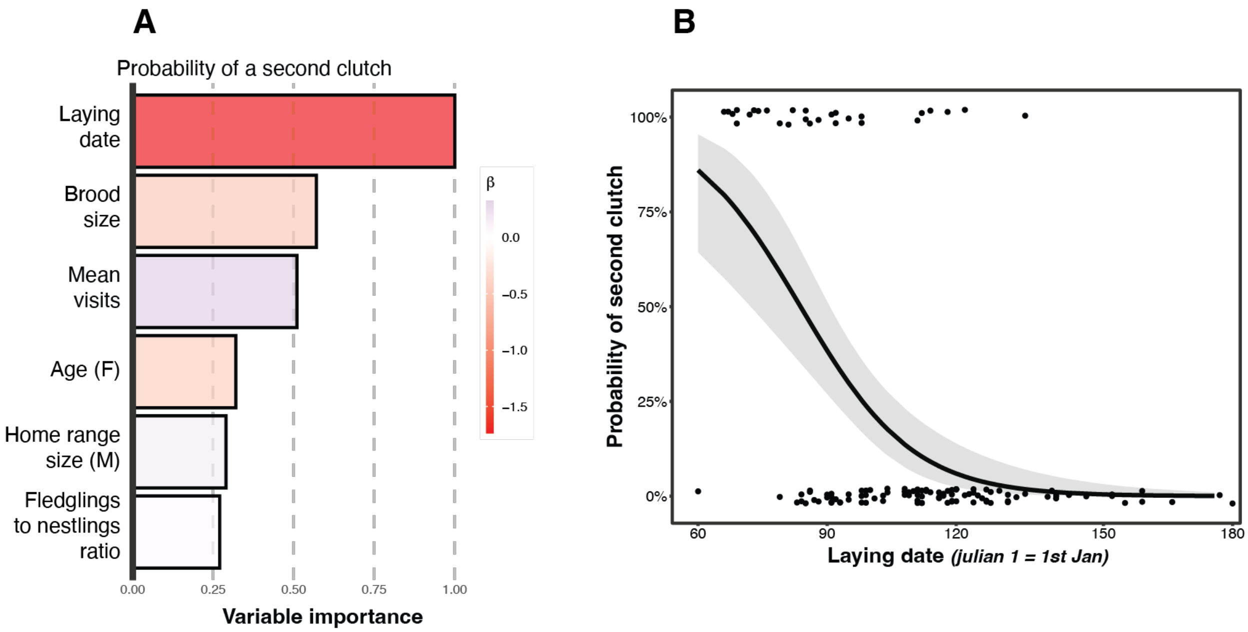


**Figure S3** – Summary of variable importance and model coefficients of averaged model (A), length of the bar is relative to variable importance (sum of Aikike weights), colour is relative to the effect direction (red = negative, blue = positive) and its intensity is proportional to the effect strength. Predicted effect of laying date (B) on the probability of re-nesting for the second time in female barn owls, grey area is relative to 95% confidence intervals.

#### Summary of univariate regression models represented in Fig. 5A

Annual sum of eggs per female (Negative binomial distr.)

| *Predictors* | *Incidence Rate Ratios* | *Conf.Int. (2.5%-97.5%)* | *Random eff. $\sigma^2_{Season}$* | *N* | *AICc* | *$R^2_{Marginal}$* |
|---|---|---|---|---|---|---|
| n. nest sites visited | 1.093** | 1.033 – 1.157 | 0.14 | 171 | 850.72 | 0.046 |
| median time at nest sites | 1.062* | 1.003 – 1.124 | 0.15 | 171 | 855.61 | 0.021 |
| prospecting[0,1] [1] | 1.131 | 0.996 – 1.285 | 0.15 | 171 | 856.06 | 0.020 |
| Mean visits | 1.069* | 1.009 – 1.132 | 0.15 | 171 | 854.69 | 0.026 |

* Null hypothesis value outside the confidence interval
* *p<0.05* ** *p<0.01* *** *p<0.001*

**Table S14** – Summary table of four univariate models having the annual sum of eggs laid per female as response variable. This table highlights the predictive difference between predictors through AICc, showing Odds Ratios and 95% C.I..

#### Summary of univariate regression models represented in Fig. 5D

**Annual sum of fledglings per female** (Poisson distr.)

| *Predictors* | *Incidence Rate Ratios* | *Conf.Int. (2.5%-97.5%)* | *N* | *AICc* | $R^2_{Marginal}$ |
|---|---|---|---|---|---|
| n. nest sites visited | 1.041 | 0.970 – 1.116 | 171 | 712.55 | 0.008 |
| median time at nest sites | 0.993 | 0.924 – 1.069 | 171 | 713.75 | 2.049e-04 |
| prospecting[0,1] [1] | 1.006 | 0.864 – 1.170 | 171 | 713.77 | 3.498e-05 |
| Mean visits | 1.003 | 0.933 – 1.078 | 171 | 713.77 | 4.953e-05 |

* Null hypothesis value outside the confidence interval
* *p<0.05* ** *p<0.01* *** *p<0.001*

**Table S15** – Summary table of four univariate models having the annual sum of fledglings per female as response variable. This table highlights the predictive difference between predictors through AICc, showing Odds Ratios and 95% C.I..
